# Gene and protein sequence features augment HLA class I ligand predictions

**DOI:** 10.1101/2023.09.03.556079

**Authors:** Kaspar Bresser, Benoit P Nicolet, Anita Jeko, Wei Wu, Fabricio Loayza-Puch, Reuven Agami, Albert JR Heck, Monika C Wolkers, Ton N Schumacher

## Abstract

The sensitivity of malignant tissues to T cell-based cancer immunotherapies is dependent on the presence of targetable HLA class I ligands on the tumor cell surface. Peptide intrinsic factors, such as HLA class I affinity, likelihood of proteasomal processing, and transport into the ER lumen have all been established as determinants of HLA ligand presentation. However, the role of sequence features at the gene and protein level as determinants of epitope presentation has not been systematically evaluated. To address this, we performed HLA ligandome mass spectrometry on patient-derived melanoma lines and used this data-set to evaluate the contribution of 7,124 gene and protein sequence features to HLA sampling. This analysis reveals that a number of predicted modifiers of mRNA and protein abundance and turn-over, including predicted mRNA methylation and protein ubiquitination sites, inform on the presence of HLA ligands. Importantly, integration of gene and protein sequence features into a machine learning approach augments HLA ligand predictions to a comparable degree as predictive models that include experimental measures of gene expression. Our study highlights the value of gene and protein features to HLA ligand predictions.

## Introduction

Spontaneous or immunotherapy-induced recognition and destruction of malignant tissues by the T cell-based immune system is, to a large extent, dependent on presentation of HLA class I bound peptides to antigen-specific CD8 ^+^ T cells^1–3^. Consequently, the composition of the pool of peptide-HLA class I complexes at the cell surface—or the HLA class I ligandome—strongly determines the ‘visibility’ of tumor cells to CD8 ^+^ cytotoxic T cells. Understanding the various factors that define the composition of this HLA ligandome is thus of major value for cancer immunotherapy.

The HLA class I ligandome is primarily generated through the intracellular degradation of proteins by the proteasome, and subsequent translocation of peptide fragments into the ER lumen by the transporter associated with antigen processing (TAP). These peptides can undergo further trimming by ER-resident aminopeptidases, bind to the peptide-binding groove of HLA class I molecules, and finally traffic to the cell surface to be presented to the immune system^4,5^. The number of peptides that can theoretically be generated from the human proteome is vast, adding up to approximately 10 ^7^ distinct peptides for 9 - meric species alone^6^. This large space poses a substantial challenge in the prediction of the HLA ligandome of a cell population of interest. Over the past decades, significant advances have been made in reducing this complexity, primarily by focusing on characteristics of the peptide itself or its surrounding sequence. Specifically, HLA class I ligands bind to the peptide-binding groove of HLA class I through shared ‘anchor’ residues, a feature that has been leveraged in the development of predictive algorithms^7,8^. In addition, the predictable cleavage preference of the proteasome ^9^ has been used to improve epitope prediction accuracy ^10,11^.

Beyond local sequence characteristics, a number of protein-level features are expected to play an important role in the generation of HLA binding peptides, for instance by tuning protein abundance and turn-over ^12–14^ . In prior work, transcriptome measurements have been used as a proxy for protein expression to aid HLA ligand predictions. However, mRNA and protein abundance correlate poorly in most mammalian c^1^e^5–^l^1^l^7^s, primarily due to post-transcriptional regulation. Such post-transcriptional regulation includes the activity of RNA-binding proteins and non-coding RNA species, and sequence intrinsic features (e.g. GC content and codon usage), which can affect the translational output of mRNAs^18,19^ . Furthermore, post-translational modifications, including ubiquitination and glycosylation, are known to modulate protein abundance, localization, and turn-over rates^20,21^, and may thereby influence epitope sampling.

In this study, we aimed to examine the potential value of gene and protein sequence features in the prediction of the HLA class I ligands. Implementing a machine learning approach, we show that the performance of such predictions can be improved through the addition of sequence features. Importantly, predictive models that make use of such features achieve the same level of predictive power as models that incorporate experimental measurements of gene-expression levels, and the predictive value of these features was generalizable to external data. Our data exemplify how the ‘hard-coded’ information of gene and protein sequence features can be exploited to infer a cell’s proteomic content and its derivatives.

## Results

### Identification of human melanoma HLA ligandomes

To investigate putative determinants of the HLA ligandome, we performed LC-MS on pan-HLA immunoprecipitates of three melanoma lines (**Fig. 1A**), resulting in the identification of 18,819 peptides derived from 6,286 proteins at a false discovery rate of <1%. The length distribution of the LC-MS detected peptides closely matched that of known melanoma-derived HLA ligands (IEDB ^22^, **Fig. 1B**), with the vast majority of peptides consisting of 9-meric species. Examination of positional frequencies of each amino acid revealed strong usage biases at position 2 and 9 (**Fig. 1C-D**). To assess whether this observed amino acid enrichment was explained by the known ligand preference of the HLA class I haplotypes expressed by these tumor lines, 9-meric peptide sequences from each melanoma line were clustered using the GibbsCluster algorithm ^23^. This analysis revealed dominant motifs present in each of the HLA ligandomes that closely matched the corresponding HLA haplotype consensus binding motifs for 11/11 HLA A and B alleles and 5/6 HLA C alleles (**Fig. 1E, Supplementary Figure 1**). In addition, HLA class I binding affinity predictions showed that the majority of LC-MS detected peptides (61.5–91.2%) were predicted to form ligands for at least 1 of the expressed HLA alleles (**Fig. 1F, Supplementary Figure 1**).

**Figure 1.**
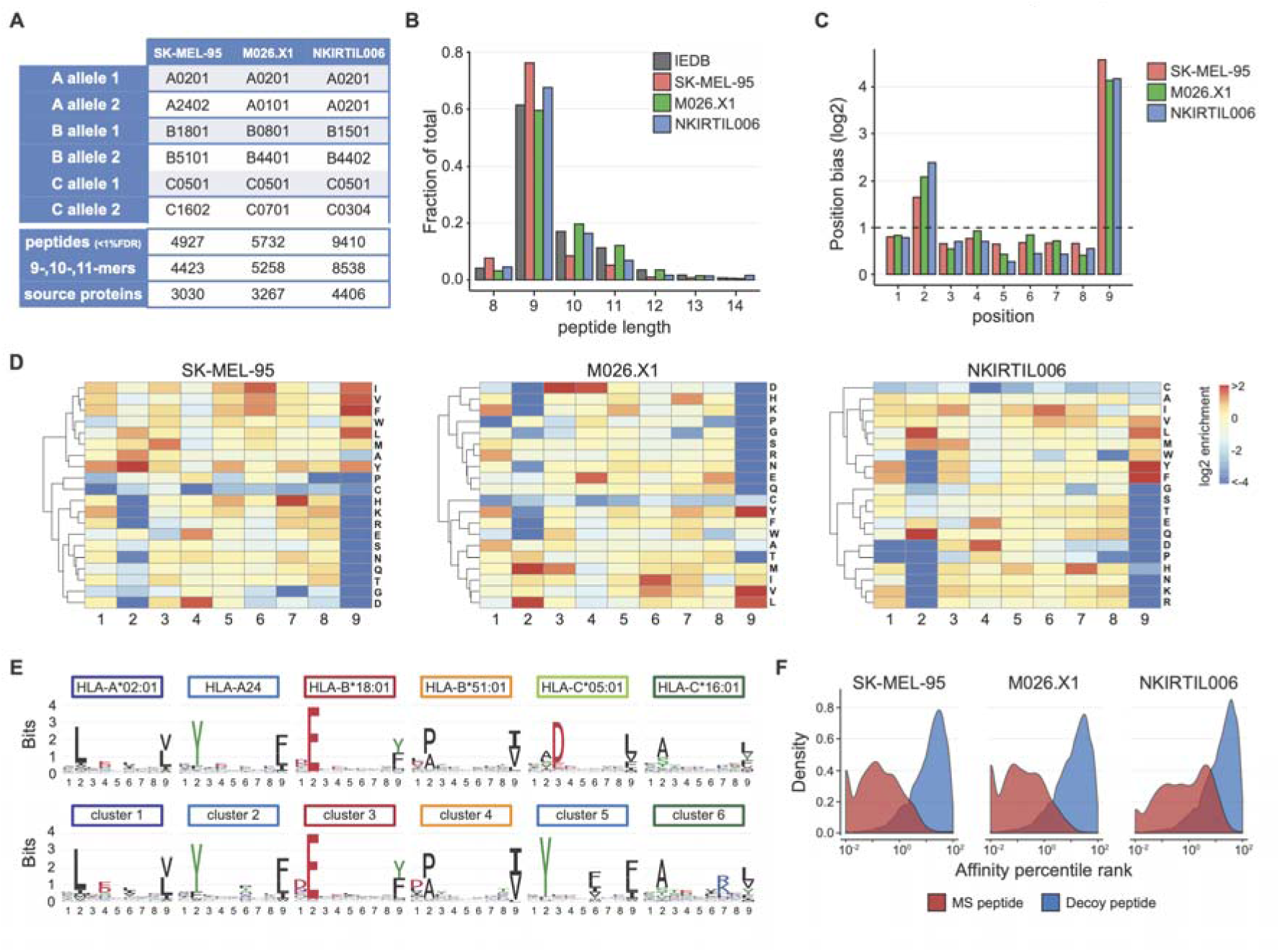
Identification of HLA ligandomes. (**A**) HLA class I haplotype of the melanoma lines used, and number of peptides and source proteins identified. (**B**) Peptide length distribution of each LC-MS dataset, compared to the peptide length distribution of known melanoma-derived HLA class I ligands deposited to IEDB. (**C-D**) Enrichment of indicated amino acids, relative to amino acid occurrence in the proteome, at each position of all 9-meric species in the datasets. Summary depicting the median of the absolute enrichment values of all amino acids for each position (C), and heatmaps visualizing hierarchical clustering of amino acid enrichment (E) are shown. (**G**) Sequence logos of all 9-mer ligands deposited to IEDB for the HLA class I alleles expressed by SK-MEL-95 (top), and the sequence logos of 6 peptide clusters obtained using the GibbsCluster algorithm (bottom). The number of clusters was constrained to the number of expressed HLA class I alleles. (**H**) Affinity percentile rank scores of LC-MS detected peptides, compared to randomly drawn peptides from transcribed genes of each melanoma line (decoy peptides).

### Gene and protein features inform on HLA sampling

Gene and protein sequence features, such as post-transcriptional or post-translational modification sites, have been shown to provide information on mRNA or protein abundance ^19,24–26^. To determine whether such features can be employed to predict the presence of HLA ligands within the proteome, we made use of a library of 7,124 sequence features. This feature library includes codon and amino acid usage, RNA-binding motifs from 142 RNA-binding proteins (RBPs), predicted miRNA (miR) binding scores, and RNA modification sites that were separately identified in the 5’ UTR, 3’ UTR and coding sequence ^27^. Predicted post-translational modification (PTM) sites, such as ubiquitination, acetylation, and malonylation sites were additionally included. Of note, this sequence feature library comprises predicted mRNA and protein modification sites, rather than any experimental measurement of such modifications in the cell systems used.

To assess whether individual sequence features can inform of HLA sampling, we first explored the contribution of a subset of features that could be assigned to five major feature classes (5’ UTR, CDS, 3’ UTR, miR binding, PTM) and that displayed a substantial degree of variance across the proteome (**Fig. 2A**, 5,771 out of 7,124 features in the library). A set of 2,000 HLA ligands was drawn from each tumor line and supplemented with a 2-fold excess of decoy peptides that were randomly sampled from the transcribed genome. This dataset was then used to train individual Random Forest classifiers for each tumor line and each sequence feature class, which were subsequently used to determine the importance of these sequence features to each of the obtained classifiers (**Fig. 2B, Supplementary Figure 2A**, showing normalized importance plots and RF metrics). The importance of sequence features was highly consistent between the different melanoma ligandome datasets, indicating that a shared set of features reliably informed on the presence of HLA ligands (**Fig. 3C**). Furthermore, direct comparison of the occurrence of high-importance sequence features within source proteins of HLA ligands and decoy peptides revealed significant differences for a set of sequence features (**Supplementary Figure 2**). For example, HLA ligands were preferentially sampled from proteins that contained a higher number of predicted sites for ubiquitination and acetylation, two PTMs that can regulate targeted proteasomal degradation and protein stability^28–30^ (**Fig. 2D**). Predicted N1-methyladenosine (m1A) sites within the 5’ UTR were also enriched in the mRNA of source proteins of HLA ligands, an effect that appears consistent with the prior observation of enhanced translation efficiency of m1A-modified mRNA molecules^25^. In contrast, 5’UTR length and occurrence of G-rich motifs in the CDS, features that have previously been suggested to negatively impact mRNA levels and translation, respectively ^31,32^, were negatively associated with the presence of HLA ligands (**Fig. 2D**).

**Figure 2.**
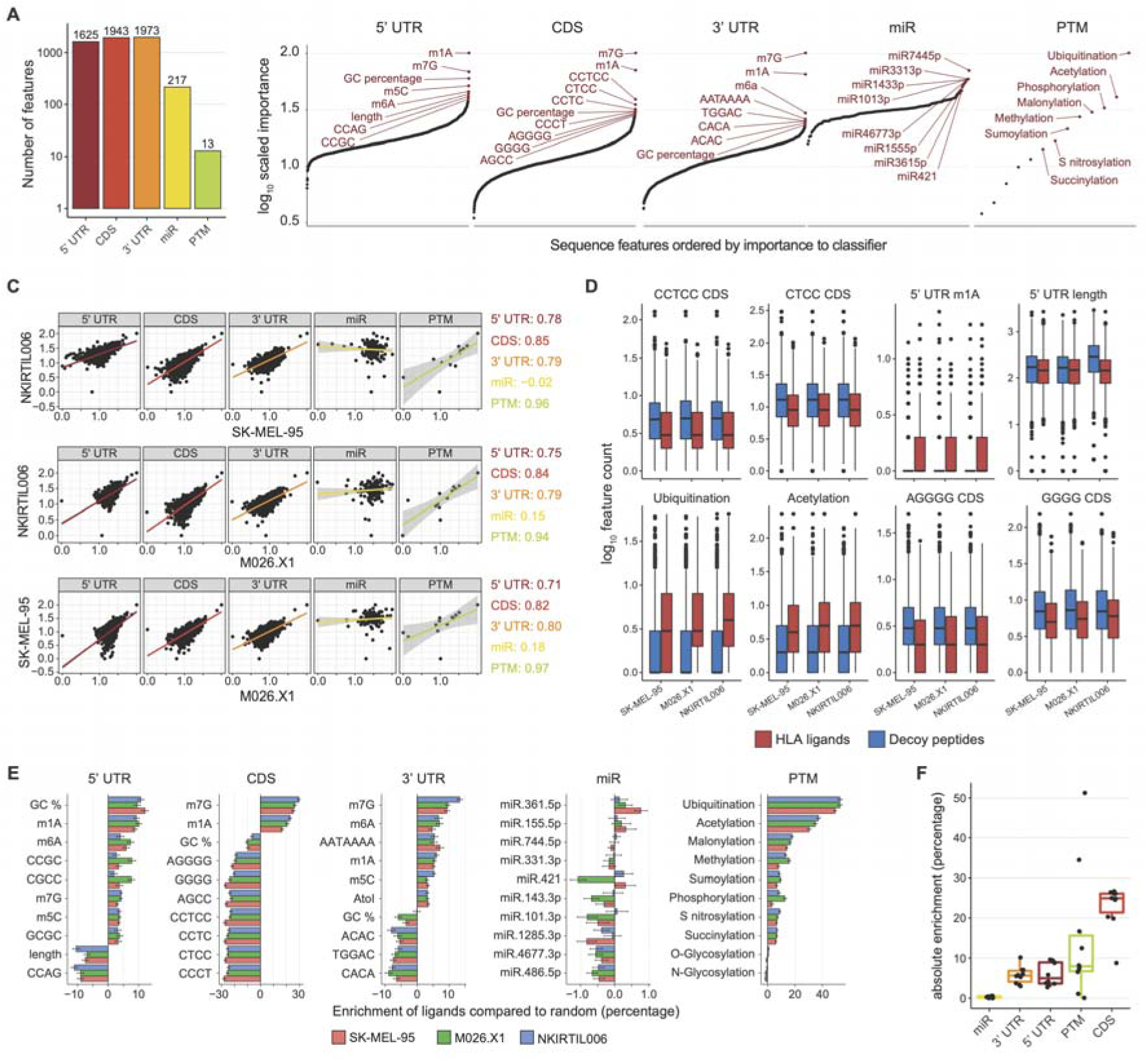
Sequence features inform on HLA sampling. Random forest models were trained using HLA ligandome data from each melanoma line and using individual classes of gene and protein sequence features to identify HLA ligands. (**A**) Sequence feature classes used to fit random forest classifiers for each melanoma line. Values indicate number of features per class. (**B**) Mean importance of sequence features to the models of each class. Feature importance represents the mean decrease in accuracy when that sequence feature is not included in the model, all importance scores are re-scaled per random forest model to a 0-100 scale. Dots indicate individual features. (**C**) Comparison of the importance of all sequence features across the individual random forest models. Dots indicate individual features, linear regressions are shown as colored lines, and 95% confidence intervals as greyed areas. Colored text denotes the respective Pearson correlation coefficients. (**D**) Comparison of sequence feature occurrence between 500 LC-MS detected HLA ligands and the same number of decoy peptides. Selected sequence features are shown. Boxplots indicate group median and 25th and 75th percentiles, whiskers indicate the interquartile range multiplied by 1.5, and dots signify individual peptides. (**E**) HLA ligands and decoy peptides were either ranked at random or by the indicated sequence feature, and the number of HLA ligands in the top 50% ranked peptides was quantified. Data depict the relative increase in HLA ligands as compared to random, bars indicate the mean percentage increase of 50 bootstraps, error bars depict 95% confidence intervals. (**F**) Comparison of averaged absolute enrichment values between feature classes. Boxplots indicate group median and 25th and 75th percentiles, whiskers indicate the interquartile range multiplied by 1.5, and dots signify individual features.

**Figure 3.**
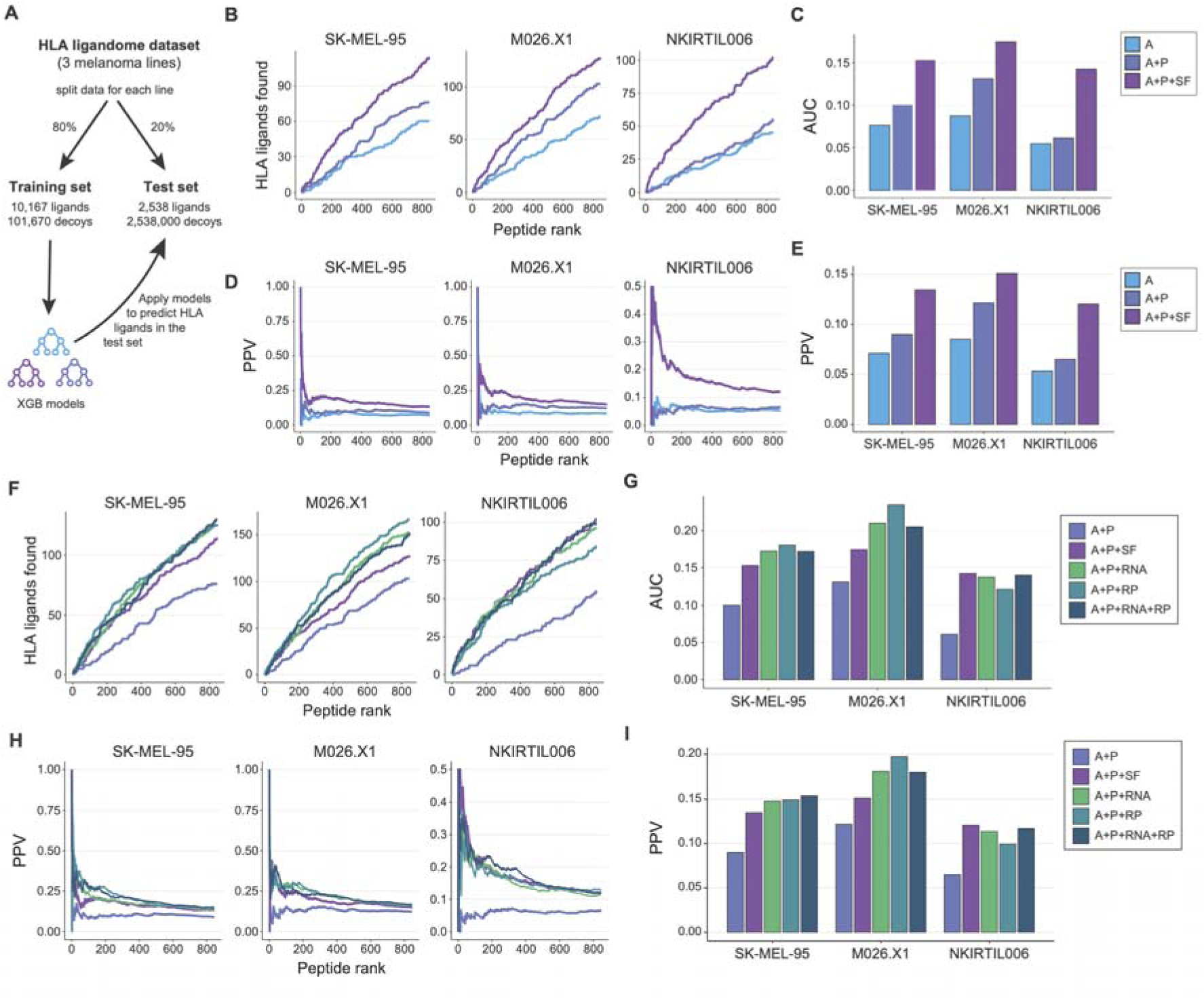
Value of sequence features in HLA ligand predictions. (**A**) The melanoma line data set was split into a training set and test set at an 80/20 ratio. The training set was used to build XGB classifiers using different combinations of features. (**B-C**) Number of true HLA ligands observed in the top 0.1% of predicted peptides from the matched melanoma line test set by each of the indicated models. Line graphs depicting the cumulative sum **B**() and bar charts depicting AUCs (**C**) are shown. (**D-E**) Positive predictive value (PPV) at each peptide rank within the top 0.1% of predicted peptides from the melanoma line test set by each of the indicated models. (**F-G**) Quantity of true HLA ligands observed in the top 0.1% of predicted peptides from the melanoma line test set by each of the indicated models. Line graphs depicting the cumulative sum (**F**) and bar charts depicting AUCs (**G**) are shown. (**H-I**) Positive predictive value (PPV) at each peptide rank within the top 0.1% of predicted peptides from the melanoma line test set by each of the indicated models. Features used to build classifiers were predicted HLA class I affinity (A), predicted proteasomal processing (P), transcript abundance (RNA), ribosome occupancy (RP), and the sequence feature library (SF).

To understand the ability of individual sequence features to contribute to HLA ligand prediction in a more quantitative manner, a custom enrichment score was calculated for each of the selected features (see methods). In brief, the set of HLA ligands and decoy peptides was sorted by the occurrence of each feature or was arranged in a random manner. Subsequently, the quantity of HLA ligands present in the top 50% ranked peptides was compared between these two cases, reflecting the benefit of each feature when used as a single determinant. In concordance with the prior analysis (**Fig. 2C**), miR binding site quantities exhibited no detectable bias toward HLA ligands or decoy peptides. In contrast, sequence features from the other classes showed a consistent capacity to enrich or deplete for the presence of HLA ligands (**Fig. 2E**). The most prominent associations were observed in the CDS and PTM classes (**Fig. 2F**), with some features increasing the number of ligands detected by more than 20%. Notably, computed m1A and N7-methylguanosine (m7G) sites were predictive of the presence of HLA ligands in the protein product irrespective of their location within the coding sequence or the untranslated regions (**Fig. 2E**), an observation that aligns with their general translation-enhancing capacity^25,33^ . Intriguingly, even though GC content was consistently informative on HLA sampling, its directionality was context dependent (positively correlated in the 5’ UTR and negatively correlated in the 3’UTR and CDS), in line with prior reports suggesting that GC content may influence mRNA levels differently depending on location^6,17,34^. Together, the above analyses show that gene and protein sequence features can individually inform on the presence of HLA ligands.

### Sequence features augment HLA ligand predictions

Having shown that individual sequence features can inform on HLA sampling, we next assessed whether these features can be leveraged to improve HLA ligand prediction models. To this end, the melanoma HLA class I ligand dataset was divided into a training set (80%) and test (20%) set that were supplemented with a 4-fold and 1,000-fold excess of decoy peptides, respectively. To evaluate the added value of sequence features to classical HLA ligand prediction methods, such as netMHC (HLA affinity) and netChop (proteasomal processing), the training set was used to generate multiple XGBoost^35^ classifier models (**Fig. 3A**), each integrating a different set of explanatory variables. As reported previously ^8,11^, both computed HLA affinity and proteasomal processing were strongly predictive of HLA sampling (**Supplementary Figure 2C, D**). Importantly, applying the obtained models to predict HLA ligands in the test set revealed that the classifier that included sequence feature information consistently and substantially outperformed the models that lacked this information. Specifically, the model including sequence features consistently ranked true HLA ligands at a higher position (**Fig. 3B-C**) and increased the positive predictive value by approximately 1.5-fold (**Fig. 3D-E**).

To determine whether the sequence features that were highly informative of HLA sampling when testing separate feature classes (figure 2) were also substantially contributing to the XGBoost classifier, the importance of those features was examined. This assessment revealed that the top scoring features in the prior analyses, such as predicted ubiquitination, acetylation, m7G, and m1A sites, were likewise dominant contributing factors in the XGBoost classifier (**Supplementary Figure 3D**). Furthermore, after predicted affinity and proteasomal processing, PTM and CDS features were generally assigned the highest importance scores (**Supplementary Figure 3E**), underlining their significance in HLA ligand prediction.

### Sequence features can match ‘wet lab’ measures of gene transcription and translation

mRNA abundance measurements have been applied in prior studies to enhance the quality of HLA ligand prediction classifiers^36,37^, and a number of recent efforts have established analysis of ribosome occupancy as a measure of active protein translation^38,39^. To understand the value of protein and gene sequence features relative to these experimental measurements of either gene expression or ribosome occupancy, we generated mRNA sequencing (transcript abundance) and ribosome profiling (protein translation activity) datasets for each of the melanoma lines. Consistent with previous reports, both measures of gene expression were strongly indicative of HLA sampling and the combination of transcript level data and ribosome occupancy data offered little additional benefit (**Supplementary Figure 3A-E**). Next, to directly compare the predictive value of sequence features versus these ‘wet-lab’ measures of gene transcription and protein translation, additional XGBoost classifiers were trained that included these metrics. Application of this new set of models on the test dataset revealed that the XGBoost model that included sequence features was able to predict true HLA ligands at an equal potency as models that incorporated wet-lab measurements of gene transcription and protein translation (**Fig. 3F-I**). Interestingly, addition of wet-lab measurements of gene transcription and protein translation in a model that contained sequence features did not consistently improve predictiveness (**Supplementary Figure 3F-G**), indicating that the predictive value of sequence features and wet-lab measurements is largely redundant. Together, these data show that gene and protein sequence features jointly provide a similar degree of information on HLA ligandome composition as experimentally obtained expression levels.

### Sequence feature-based HLA ligand classifiers are generalizable to external data

To understand whether the value of sequence features in HLA ligandome prediction is generalizable, we subsequently tested whether the above observations could be extended to another tumor type, other HLA class I alleles, and to an independent dataset. To this purpose, we assessed the performance of the different classifiers on an HLA ligandome data set obtained from multiple mono-allelic human B lymphoblastoid cell lines ^37^, focusing on HLA ligandome data for the 4 most common HLA class A and B alleles ^40^, plus HLA ligandome datasets for 4 common HLA A and B alleles that were absent from the original melanoma dataset. From each of these ligandomes, 350 9-meric HLA ligands were sampled and supplemented with a 1,000-fold excess of decoy peptides, resulting in 8 individual test datasets. Application of the classifiers trained on internal ligandome data to these external test sets showed that models that included sequence features outperformed predicted HLA affinity- and processing-based models in 7 out of 8 datasets (**Fig. 4A-D**). Furthermore, the performance of the sequence feature-based classifier was equal to that of XGBoost models that included the matched gene expression information from the external data sets (**Fig. 4F-G**), demonstrating that the value of gene and protein feature-based prediction models is generalizable across cell types and datasets.

**Figure 4.**
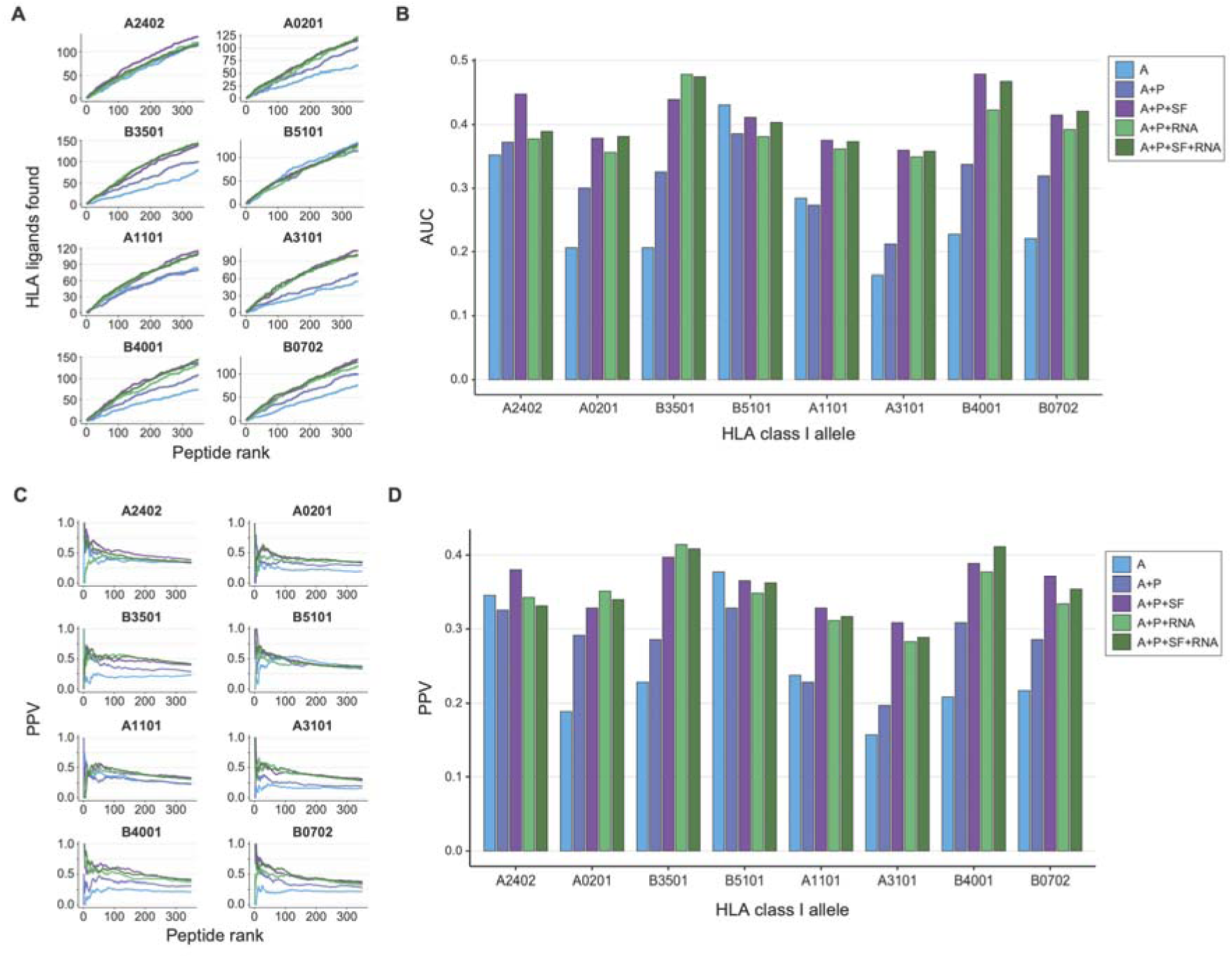
Sequence feature-based XGBoost models are generalizable to external data. XGBoost classifiers were validated using HLA ligandome data from 6 mono-allelic cell lines from a previously published dataset^36^. From each cell line, 350 true HLA ligands were supplemented with 350,000 decoy peptides. (**A**) Cumulative sum graphs of true HLA ligands observed in the top 0.1% of predicted peptides, by each of the indicated models. (**B**) AUC values for each of the graphs shown in panel A. (**C**) Positive predictive value (PPV) at each peptide rank within the top 0.1% of predicted peptides by each of the indicated models. (**D**) PPV within the top 0.1% of predicted peptides by each of the indicated models. Features used to build classifiers were predicted HLA class I affinity (A), predicted proteasomal processing (P), transcript abundance (RNA), and the sequence feature library (SF).

## Discussion

Gene and protein sequence features represent a class of ‘hard-coded’ regulators of protein expression, influencing this process at many different levels. In this study, we leveraged a large set of such gene and protein features to assess their contribution to the composition of the HLA ligandome. Through this effort, we demonstrate that sequence features can augment HLA ligand predictions, and that the predictive gain obtained in these models is equal to that of models that incorporate experimentally obtained gene expression and protein translation data.

While not formally assessed here, it is expected that at least some of the sequence features contribute to HLA ligand predictions by providing a proxy for protein abundance. This notion is supported by the observation that sequence features such as mRNA region length, GC content, and post-transcriptional modifications can be used to help predict protein levels^17^. Furthermore, predicted RNA methylation sites were among the features that were most prominently associated with the presence of HLA ligands, an observation that may be explained by their known modulatory effect on both mRNA stability and translation efficiency ^25,26,33,41^. In addition to features involved in mRNA regulation and translation, our data reveal that the predicted occurrence of several PTM sites informed on the presence of HLA ligands. For ubiquitination, the PTM that displayed the highest predictive value, its positive association with HLA ligand yield may be caused by an enhanced accessibility to proteasomal degradation ^20,30,42^. For other PTMs that were predictive of HLA ligands, such as methylation and acetylation, their involvement in specific pathways is less well understood^21,43,44^. Our data provide correlative evidence that that these modifications influence availability of proteins to the antigen processing machinery but further work will be required to formally test this.

Improvement of HLA ligand prediction approaches remains an active field of research, with the aim to, for example, allow the more precise selection of cancer (neo)antigens for therapeutic purposes^36,37,45,46^ . Because of its generalizable nature and lack of requirement for direct transcriptome measurements, we envision that the approach described here will be of value in these efforts.

## Methods

### Patient-derived melanoma cell lines

SK-MEL-95 and M026.X1^47^ were a kind gift from Daniel Peeper (Netherlands Cancer Institute). SK-MEL-95 was originally established in the Memorial Sloan Kettering Cancer Center. NKIRTIL006 was established in house^48^.

### Cell culture

Patient-derived melanoma cell lines were cultured in RPMI (Gibco) supplemented with 8% fetal calf serum (FCS, Sigma), 100 U/ml penicillin (Gibco) and 100 µg/ml streptomycin (Gibco) at 37 °C and_2_.5% CO For mRNA sequencing and ribosome profiling, cell lines were cultured to a density of 70-90% on 150mm Corning tissue-culture treated culture dishes (Merck). For HLA ligandome LC-MS, approximately 1·10^9^ cells were cultured in Corning CELLSTACK Culture Chambers (Corning, 05-539-096).

### HLA class I peptide isolation and LC-MS/MS

HLA class I-associated peptides were isolated by isolated by immunoprecipitation of HLA class I complexes using the mouse monoclonal IgG2a antibody W6/32, as described previously ^49^. Peptides were eluted from HLA class I protein molecules using a 10% acetic acid (v/v) solution, and subsequently separated using a 10 kDa molecular weight cutoff filter. Obtained solution was then desalted into 3 fractions using in-house made c18 STAGE (STop And Go Extraction) tips, eluted with 20%, 30% and 50% acetonitrile, respectively. The resulting fractions were injected on an Agilent 1290 system using a 120-min gradient coupled to an Orbitrap Fusion mass spectrometer (Thermo Fisher Scientific). Fractions 1 and 2 were injected in triplicate, whereas fraction 3 was injected in duplicate. The LC system comprised of a 20 × 0.1 mm i.d. trapping column (Reprosil C18, 3 μm; Dr. Maisch) and a 50 × 0.005 cm i.d. analytical column (Poroshell 120 EC-C18; 2.7 μm). An LC resolving gradient of 13 to 43% Solvent B (80% acetonitrile, 20% water, 0.1% formic acid) was used. The Top Speed method was enabled for fragmentation, where the most abundant precursor ions were selected in a 3 s cycle for data-dependent EThcD. MS1 and MS2 spectra were acquired at a resolution of 60,000 (FWHM at 400 *m*/*z*) and 15,000 (FWHM at 400 *m*/*z*), respectively. RF lens voltage was set to 60. Dynamic exclusion of 18s was used. Peptide precursors of charges 2 to 6 were fragmented, if accumulated to a minimum intensity of ^5^4w·1it0hin 50 ms. In MS2, a maximum injection time of 250ms was used with a minimum intensity filter of 5·10^4^.

### HLA class I peptide analysis

RAW data files were analyzed using the Proteome Discoverer 1.4 software package (Thermo Fisher Scientific). MS/MS scans were searched against the human Swissprot reviewed database (accessed in September 2015; 20,203 entries), with no enzyme specificity using the SEQUEST HT search engine. Precursor ion and MS/MS tolerances were set to 10 ppm and 0.05 Da. Methionine oxidation was set as variable modification. The peptides-to-spectrum matches were filtered for precursor tolerance 5 ppm, <1% FDR using Percolator, XCorr >1.7, and peptide rank 1. Peptides that were between 8 and 14 amino acid long were selected for further analysis. The mass spectrometry data have been deposited to the ProteomeXchange Consortium via the PRIDE partner repository with the data set identifier PXD036277. Replicate injections displayed an overlap of approximately 70% (shared between at least 2 replicates). Consistent with their shared tissue origin, a large part of peptides detected across the melanoma lines mapped to a core group of proteins (47.6% shared between at least 2 lines). In contrast, the MS detected peptides exhibited a small degree of overlap (12.4% shared between at least 2 lines), in line with their difference in HLA haplotype.

### mRNA sequencing

Cells were cultured to an approximate density of 80%, and 1·10 ^7^ cells were subsequently dissociated using a cell-scraper in cold (4 °C) PBS, centrifuged for 10 minutes at 300x *g*, and snap-frozen in liquid nitrogen. RNA was extracted from the frozen pellets using the RNeasy Mini Kit (Qiagen). Whole transcriptome sequencing samples were prepared using the TruSeq Stranded mRNA Kit (Illumina). Single-end 65lllbp sequencing was performed on a HiSeq 2500 System (Illumina). Obtained reads were aligned to the GRCh38 reference (gencode release 21) using STAR aligner (version 2.5.2b), and transcripts were quantified using Salmon (version 0.7.0). Transcript counts belonging to a single consensus coding sequence were summed.

### Ribosome profiling

Cells were cultured to an approximate density of 80%, and 5·10 ^7^ cells were subsequently treated with 100lllμg/ml cycloheximide for 5lllminutes at 37 °C. Cells were then washed once in cold (4 °C) PBS containing 100lllμg/ml cycloheximide, dissociated using a cell-scraper in cold (4 °C) PBS supplemented with 100lllμg/ml cycloheximide, centrifuged for 10 minutes at 300x *g*, and snap-frozen in liquid nitrogen. Frozen pellets were resuspended in lysis buffer (20lllmM Tris–HCl, pH 7.8, 100lllmM KCl, 10lllmM MgCl_2_, 1% Triton X-100, 2lllmM DTT, 100lllμg/ml cycloheximide, 1× Complete protease inhibitor), and incubated on ice for 20 minutes. Lysates were sheared using a 26G needle, centrifuged for 10 minutes at 1,300x *g*, and supernatants were transferred to a clean tube. Supernatants were treated with 2lllU/μl of RNase I (Ambion) for 45lllmin at room temperature, with rotation. Next, lysates were fractionated on a linear sucrose gradient (7–47%) using the SW-41Ti rotor (Beckman Coulter) at 221,633x *g* for 2lllhours at 4 °C, without brake. Obtained sucrose gradients were then divided in 14 fractions, and fractions 7–10 (cytosolic ribosomes) were pooled and treated with PCR grade proteinase K (Roche) in 1% SDS to release ribosome protected fragments. The resulting fragments were subsequently purified using Trizol reagent (Invitrogen) and precipitated in the presence of glycogen, following the manufacturer’s instructions. For library preparation, RNA was gel-purified on a denaturing 10% polyacrylamide urea (7 M) gel. A section corresponding to 25 to 36 nucleotides—the region that comprises the majority of the ribosome-protected RNA fragments—was excised, and purified through ethanol precipitation. RNA fragments were then 3lll-dephosphorylated using T4 polynucleotide kinase (New England Biolabs) for 6 hours at 37°C in 2-(N-morpholino)ethanesulfonic acid (MES) buffer (100 mM MES-NaOH pH 5.5, 10 mM MgCl _2_, 10 mM β-mercaptoethanol, 300 mM NaCl). The 3lll adaptor was added using T4 RNA ligase 1 (New England Biolabs) for 2.5 hours at 37°C. Ligation products were 5lll-phosphorylated with T4 polynucleotide kinase for 30 minutes at 37 °C, and the 5lll adaptor was added using T4 RNA ligase 1 for 2 hours at 37 °C. Sequencing was performed on a HiSeq 2500 System (Illumina). Ribosome occupancy was calculated using the Ribomap pipeline ^50^, and was aligned to the GRCh38 reference (gencode release 21). Counts belonging to a single consensus coding sequence were summed.

### Characterization of LC-MS detected peptides

For comparison of peptide length distributions, known melanoma HLA class I ligands were downloaded from the IEDB web-interface (https://www.iedb.org) in June 2021 using the following search filters: Epitope – Any; Assay Outcome – Positive; MHC restriction – Class I; Host – Human; Disease – Melanoma.

To assess the amino acid positional biases of the LC-MS detected peptides, the dataset was filtered for 9-meric species, and the occurrence of each amino acid on each peptide position was tallied. As a reference, all expressed proteins (TPM > 0 in the mRNAseq dataset) were selected for each melanoma line, and the number occurrences of each amino acid was calculated. Amino acid enrichment was then defined as the fraction by which an amino acid occurred at a certain position divided by the fraction by which that amino acid occurred in the reference. The positional bias was defined as the median of the absolute amino acid enrichment values for each peptide position.

For binding motif analyses, 9-meric peptide sequences from each melanoma line were clustered using GibbsCluster 2.0 (command line options set to: -g 3-7 -C -D 4 -I 1 -S 5 -T -j 2 -c 1 -k 25), with the number of clusters for each melanoma line set to the number of alleles expressed by that line. Sequence logos were generated using the R package ggseqlogo. To generate reference sequence logos, all known human 9mers for each of the shown HLA class I alleles were downloaded from IEDB in June 2021.

### Peptide database construction

To investigate characteristics of HLA class I ligands, a database consisting of LC-MS detected peptides (i.e., true HLA ligands) and not-detected peptides (referred to as decoy peptides) was constructed. To this end, HLA binding scores to the HLA alleles of each melanoma line were calculated for all 9-, 10-, and 11-mers in the human proteome (GRCh38, gencode release 21) using netMHCpan 4.0. Processing scores were calculated using netChop 3.1. Separate databases were generated for each melanoma line by filtering on peptides derived from expressed proteins (TPM > 0 in the mRNAseq dataset), and assigning each peptide the highest affinity rank score out of the expressed HLA alleles. LC-MS detected peptides were then assigned as ‘true HLA ligands’ and the remainder of all peptides as ‘decoy peptides’. When this database was sampled for analyses, equal peptide length distributions were maintained between true HLA ligands and decoy peptides.

### Feature library construction

5’ UTR, coding region (CDS) and 3’ UTR nucleotide sequences were downloaded from ENSEMBL BiomaRt (release 104; accessed September 2021) for all protein-coding transcripts. RNA-binding protein motifs were acquired from ATtracT ^51^ (accessed June 2021) and filtered for human RBPs (142 RBPs; 2,271 motifs). In each transcript region (e.g., 5’ UTR, CDS, 3’ UTR), motifs were searched and counted using a custom script (see GitHub project), and GC content and nucleotide length were computed. Also included in the sequence feature library were: Codon usage (applying coRdon^52^), amino acid usage within the CDS, miR-DB ^53^ miRNA seed scores (accessed August 2021 and filtered for miRNA expressed immune cel based on previous analysis by Juzenas *et al.* ^54^), sequence homology between Human and Zebrafish (*Danio rerio*, obtained through Ensembl BiomaRt), predicted mRNA modification site occurrence per transcript region (obtained from the RMVar database^55^, accessed at https://rmvar.renlab.org/ in September 2021), and predicted post-translational modification (Acetylation, Amidation, Hydroxylation, Malonylation, Methylation, N-linked_Glycosylation, O-linked_Glycosylation, Palmitoylation, Phosphorylation, S-nitrosylation, Succinylation, Sumoylation, Ubiquitination) site occurrence (obtained from the dbPTM database ^56^, accessed at https://awi.cuhk.edu.cn/dbPTM/ in June 2021).

### Importance assessment of sequence feature classes

To assess the ability of sequence features to inform on HLA sampling, features belonging to five major classes (5’ UTR, CDS, 3’ UTR, miR binding and PTM) were extracted from the sequence feature library. The 5’ UTR, CDS, 3’ UTR classes were filtered based on their variance across the proteome using the nearZeroVar function in the caret R package (setting cutoffs at: freqRatio < 500 and percentUnique > 0.05). All putative miR binding sites and PTMs in the library were used in the analysis. The number of features left after filtering are shown in **Figure 2A** . 2,000 true HLA ligands and 4,000 decoy peptides were sampled from the peptide database of each melanoma line, and subsequently used to train individual Random Forest models for each melanoma line and each feature class to predict true HLA ligands (15 models in total). The Random Forest models were generated using the R packages randomForest and caret, using 10-fold cross validation optimizing the ROC metric. Number of trees in each forest was set to 5,000 and minimum terminal node size was set to 2. The mtry parameter was set to 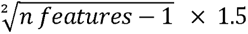. Feature importance (i.e., mean decrease in accuracy) was calculated using the varImp function from the R package caret.

Analyses examining HLA ligand enrichment potential of individual sequence features (**Figure 2D-F, Supplementary Figure 2B**) were focused on the 10 most important features in each class (defined as the highest mean importance score of the models trained for that feature class), and were performed using 3,389 true HLA ligands and 13,556 decoy peptides per tumor line. For the analysis presented in **Figure 2E-F**, a custom enrichment metric was calculated. In brief, 30% of the data was sampled and peptides were ranked either by the occurrence of a sequence feature or at random. In both cases the total number of true HLA ligands within the top 50% ranked peptides was tallied. Next, the percentage increase in true HLA ligands was calculated comparing the sequence feature ranked case versus the randomly ranked case. This process was performed for all sequence features in the analyses, and was repeated 50 times.

### XGBoost classifiers

The number of experimentally detected HLA ligands from each melanoma line was down-sampled to the number of HLA ligands in the smallest dataset to ensure each melanoma line had equal weight during the analyses. The sampled data was split into a training (80%) and a test (20%) set, and these sets were supplemented with a 4-fold or 1,000-fold excess of decoy peptides. XGBoost models were generated using the R packages xgboost and caret, using 2-times 10-fold cross validation optimizing the accuracy metric. Learning rate was set to 0.3, minimum loss reduction was set to 1.0, maximum tree depth was set to 1, sub-sampling ratio of features for each tree was set to 0.5, minimum sum of instance weight needed in a terminal leaf was set to 0.9, number of rounds was set to 1,000.

### External HLA ligandome data

Transcriptomic data was accessed from the Gene Expression Omnibus (GEO) at GSE131267 and was aligned to the GRCh38 reference (gencode release 21) using Salmon (quasi-mapping mode, version 0.7.0). Mean transcript counts were calculated between replicates, and transcripts belonging to a single consensus coding sequence were summed. HLA ligands from the Sarkizova study ^37^ were downloaded from the publisher’s website. This dataset was filtered for 9-meric peptides and peptides obtained from the mono-allelic cell lines expressing A2402, A0201, B3501, B5101, A1101, A3101, B4001 or B0702. To generate a decoy peptide pool, affinity binding ranks of all 9-meric peptides in the expressed proteome (TPM > 0) of the mono-allelic cell lines were calculated for each selected HLA allele. Processing scores were calculated using netChop 3.1. For each of the mono-allelic cell lines, 350 true HLA ligands and 350,000 decoy peptides were sampled.

## Author contributions (CRediT taxonomy)

**Kaspar Bresser:** Conceptualization, Methodology, Validation, Formal Analysis, Investigation, Data Curation, Writing – Original Draft, Writing – Review & Editing, Visualization. **Benoit P Nicolet:** Conceptualization, Methodology, Formal Analysis, Writing – Review & Editing. **Anita Jeko:** Formal Analysis, Methodology, Investigation. **Wei Wu:** Data Curation. **Fabricio Loayza-Puch:** Investigation, Methodology, Resources. **Reuven Agami:** Methodology, Resources, Supervision. **Albert JR Heck:** Conceptualization, Methodology, Resources, Supervision, Funding Acquisition. **Monika C Wolker**M**s:**ethodology, Writing – Review & Editing, Supervision **. Ton N Schumacher:** Conceptualization, Methodology, Writing – Review & Editing, Supervision, Funding Acquisition.

## Data availability

Transcriptomic and ribosome profiling data presented in this manuscript have been deposited to GEO and can be accessed under the accession number GSE211000. Mass spectrometry data have been deposited to the ProteomeXchange Consortium via the PRIDE partner repository with the data set identifier PXD036277. All statistical source data of the figures presented in this study are provided with this paper. Transcriptomic data of the Sarkizova study^37^ was accessed from GEO under the accession number GSE131267. HLA ligands from the Sarkizova study ^37^ were downloaded from the publisher’s website. Any additional data supporting the findings of this study are available from the corresponding authors upon request.

## Data availability

R scripts that were used to produce the main and extended data figures in the manuscript are available from GitHub (https://github.com/kasbress/HLA_ligandome_feature).

---

**Figure S1.**
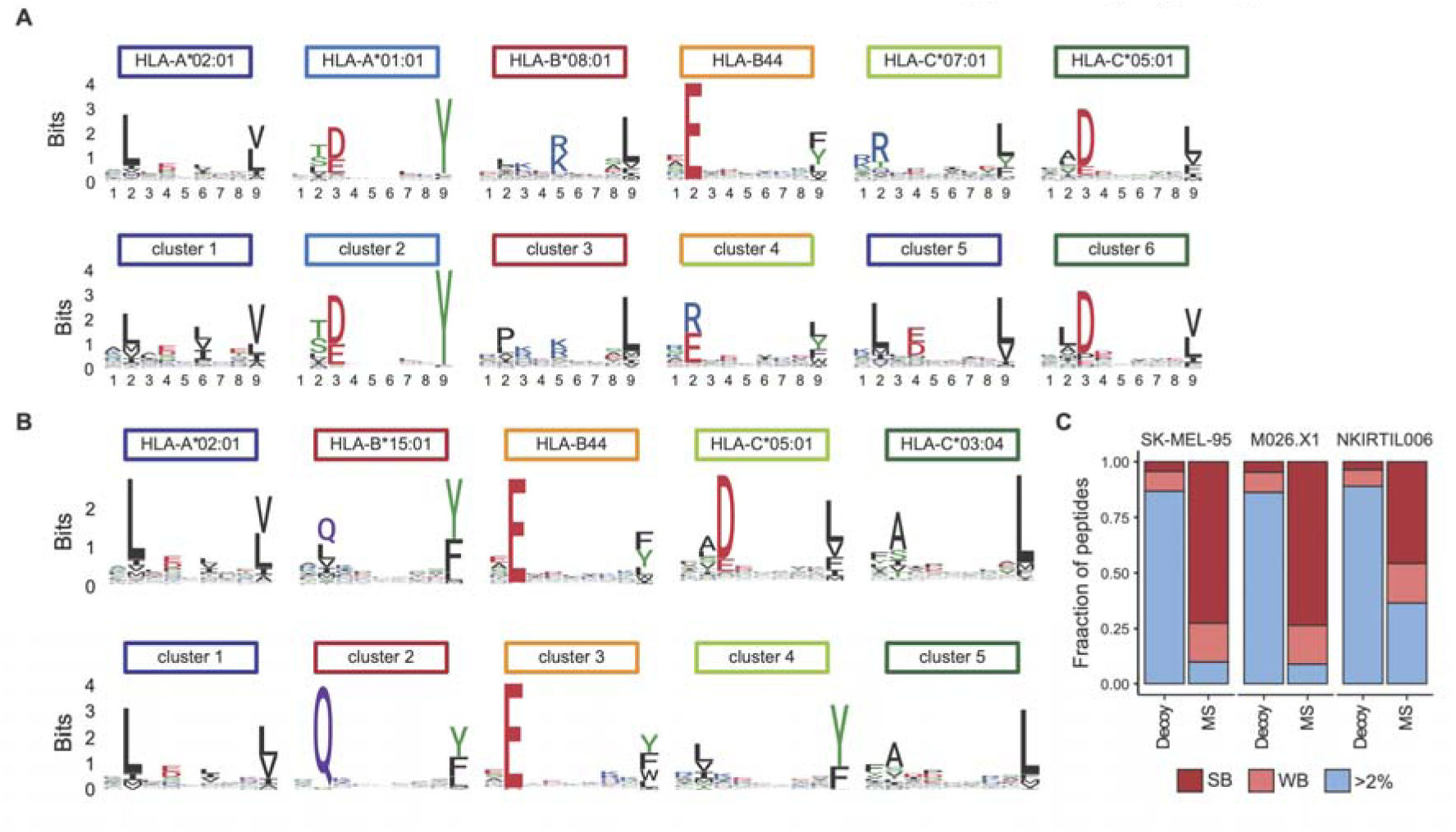
Identification of HLA class I ligandomes (related to Figure 1). (**A–B**) Sequence logos of the HLA class I alleles (top panels) expressed by M026.X1 (**A**) and NKIRTIL006 (**B**), and the sequence logos of peptide clusters obtained by the GibbsCluster algorithm (bottom panels). The number of clusters was constrained to the number of expressed HLA class I alleles. (**C**) Fraction of peptides predicted to have a <0.5 (strong binders), <2.0 (weak binders), or >2.0 percentile rank binding affinity within LC-MS detected peptides and decoy peptides, as determined by netMHCpan4-0.

**Figure S2.**
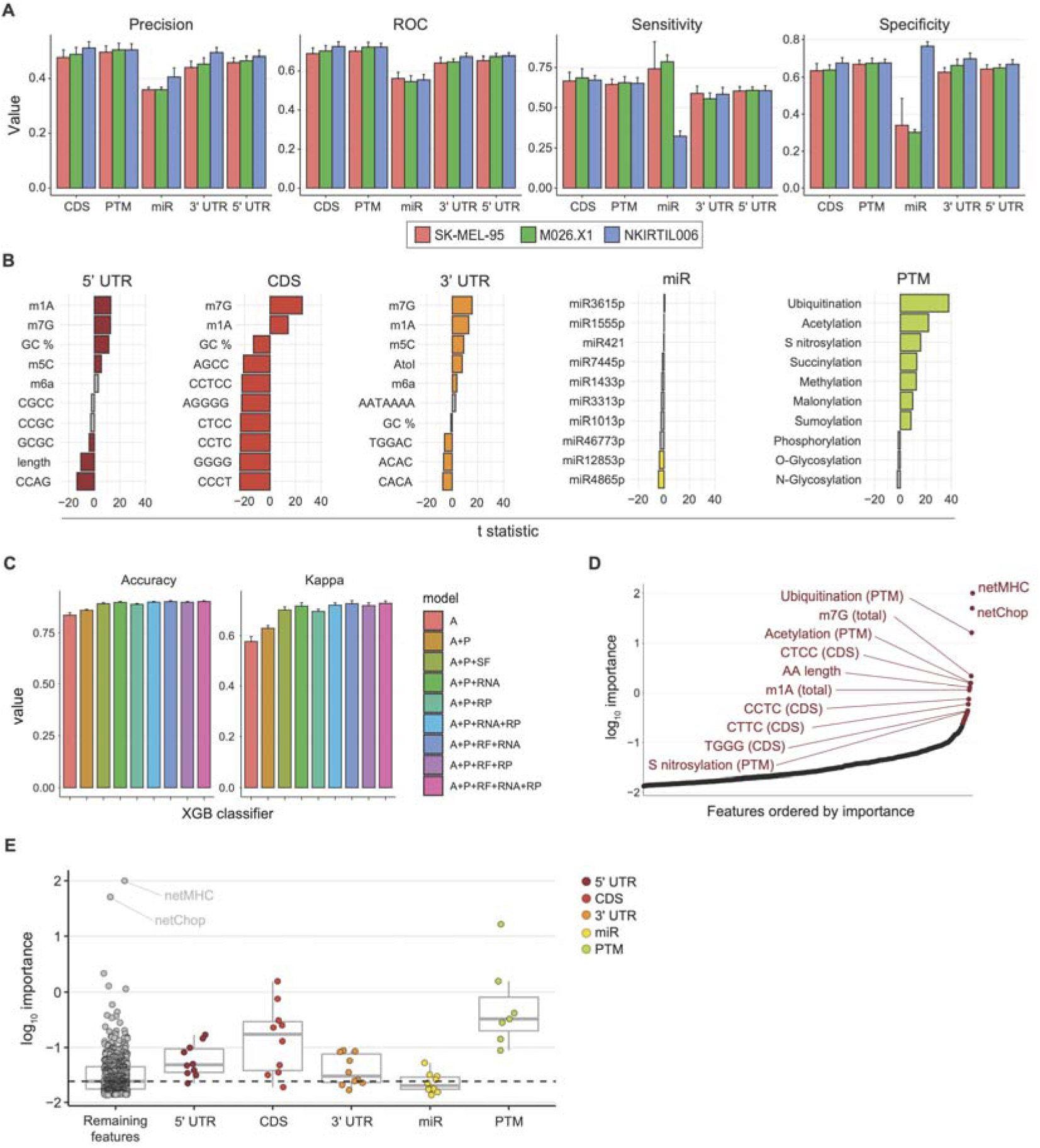
Sequence features inform on HLA sampling (related to Figure 2). (**A**) Random forest models were trained using data from each melanoma line to classify HLA ligands, using individual classes of gene and protein sequence features. Bar graphs indicate out-of-bag model performance. (**B**) t statistics obtained comparing counts of indicated sequence features between LC-MS detected and decoy peptides. Top 10 features with the highest importance to each random forest model are shown. (**C**) Out-of-bag model performance of XGBoost classifiers generated in Figure 3. (**D**) Importance of features to the A+P+SF XGBoost model. The top 12 features are highlighted. (**E**) Feature importance of the top 10 features in each class identified in Figure 2, compared to the feature importance of the remaining features in the library. Dashed line indicated the median importance of all features in the A+P+SF XGBoost model. Boxplots indicate group median and 25th and 75th percentiles, whiskers indicate the interquartile range multiplied by 1.5, and dots signify individual features.

**Figure S3.**
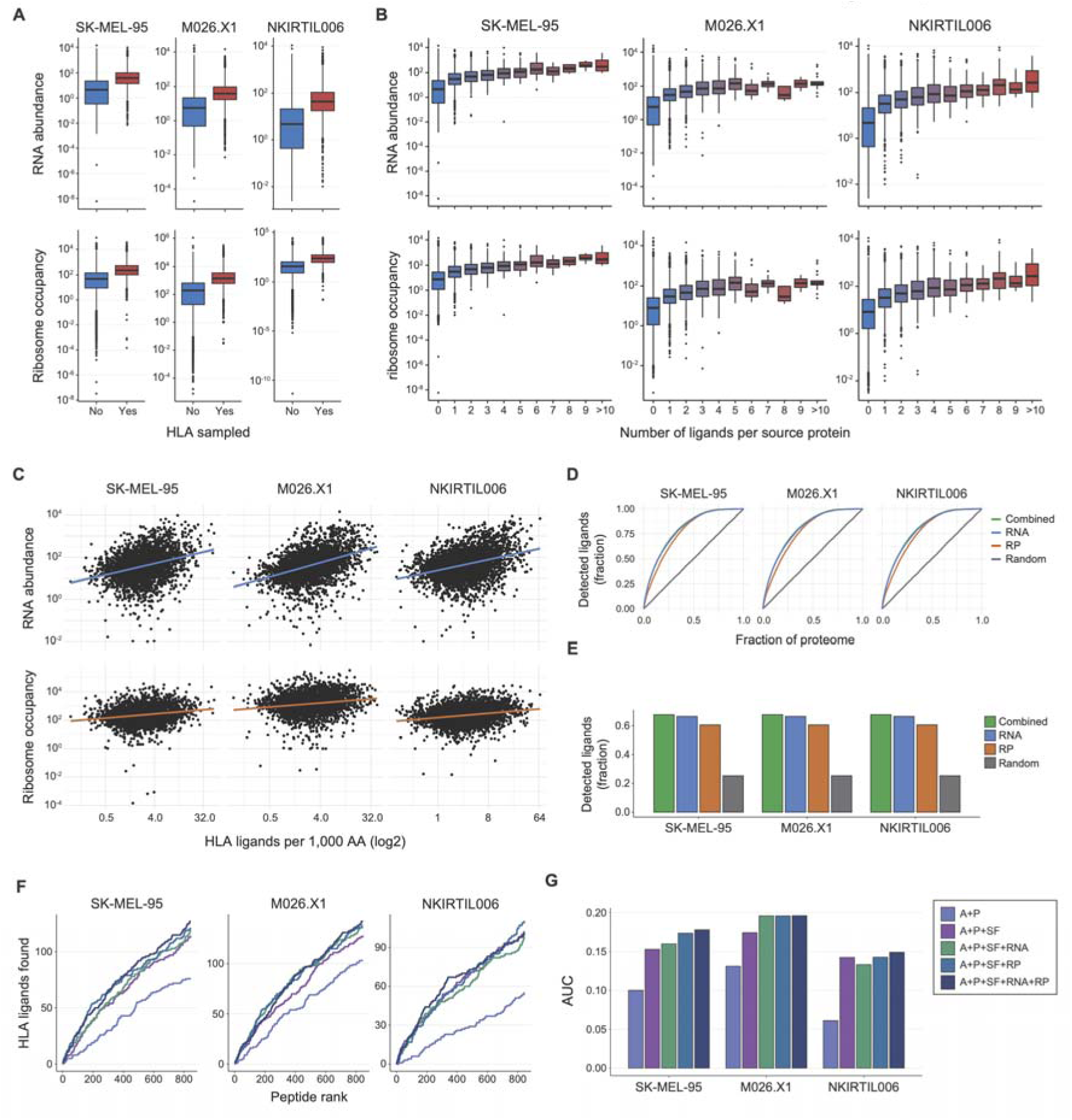
Association of HLA ligandome composition with RNA abundance and ribosome occupancy. (**A**) RNA abundance (top) and ribosome occupancy (bottom) of proteins for which either HLA ligands were or were not detected by LC-MS. (**B**) RNA abundance (top) and ribosome occupancy (bottom) of source proteins, binned by the number of HLA ligands detected by LC-MS. (**C**) HLA sampling density of each source protein, calculated as the number of detected HLA ligands per 1,000 amino acids, plotted against their respective RNA abundance (top) and ribosome occupancy (bottom). (**D-E**) Predictiveness of the assessed expression metrics. Source proteins were ranked by either RNA abundance, ribosome occupancy or by a randomly generated metric (obtained by shuffling RNA abundance data). In addition, a combined ranking was obtained by averaging the rankings of RNA abundance and ribosome occupancy metrics. Line plots (**D**) depict the fraction of detected HLA ligands from that melanoma line as a function of the fraction of the analyzed proteome (cumulative protein length). Bar charts (**E**) depict the fraction of HLA ligands observed within the top quartile of the proteome (cumulative protein length). (**F-G**) Number of HLA ligands observed in the top 0.1% of predicted peptides from the melanoma line test sets by each of the indicated models. Line graphs depicting the cumulative sum (**F**) and bar charts depicting AUCs (**G**) are shown.

